# Super-Mendelian inheritance mediated by CRISPR/Cas9 in the female mouse germline

**DOI:** 10.1101/362558

**Authors:** Hannah A. Grunwald, Valentino M. Gantz, Gunnar Poplawski, Xiang-ru S. Xu, Ethan Bier, Kimberly L. Cooper

**Affiliations:** Division of Biological Sciences, Section of Cellular and Developmental Biology, University of California San Diego, La Jolla, California 92093, USA; Department of Neurosciences, University of California San Diego, La Jolla, California 92093, USA; Tata Institute for Genetics and Society, University of California San Diego, La Jolla, California 92093, USA

**Author notes:** Equal contributions (Hannah A. Grunwald, Valentino M. Gantz, Gunnar Poplawski). Current address: Department of Medicine, National University of Singapore, Singapore 119228, Republic of Singapore (Gunnar Poplawski).

## Abstract

A gene drive biases the transmission of a particular allele of a gene such that it is inherited at a greater frequency than by random assortment. Recently, a highly efficient gene drive was developed in insects, which leverages the sequence-targeted DNA cleavage activity of CRISPR/Cas9 and endogenous homology directed repair mechanisms to convert heterozygous genotypes to homozygosity. If implemented in laboratory rodents, this powerful system would enable the rapid assembly of genotypes that involve multiple genes (e.g., to model multigenic human diseases). Such complex genetic models are currently precluded by time, cost, and a requirement for a large number of animals to obtain a few individuals of the desired genotype. However, the efficiency of a CRISPR/Cas9 gene drive system in mammals has not yet been determined. Here, we utilize an active genetic “CopyCat” element embedded in the mouse *Tyrosinase* gene to detect genotype conversions after Cas9 activity in the embryo and in the germline. Although Cas9 efficiently induces double strand DNA breaks in the early embryo and is therefore highly mutagenic, these breaks are not resolved by homology directed repair. However, when *Cas9* expression is limited to the developing female germline, resulting double strand breaks are resolved by homology directed repair that copies the CopyCat allele from the donor to the receiver chromosome and leads to its super-Mendelian inheritance. These results demonstrate that the CRISPR/Cas9 gene drive mechanism can be implemented to simplify complex genetic crosses in laboratory mice and also contribute valuable data to the ongoing debate about applications to combat invasive rodent populations in island communities.

## Main Text

A cross of mice that are heterozygous for each of three unlinked genes must produce 146 offspring for a 90% probability that one will be triple homozygous. The likelihood decreases further if any of the three genes are close together, or genetically linked, on the same chromosome. The cost, time, and animal requirements are therefore prohibitive for certain complex genotypes that might model multigenic human diseases, permit assembly of multiple optogenetic elements to study nervous system function, or recapitulate the phenotypes of evolutionarily divergent species using laboratory mice.

Recently, CRISPR/Cas9-mediated gene drive systems were developed in insects^1–3^, which could surmount the obstacles to assembling complex genotypes in laboratory rodents by increasing the frequency of inheritance of desired alleles. These gene drives utilized self-propagating elements, which we refer to broadly as “active genetic elements”, that can carry transgenes or delete and replace sequences with orthologous sequences from other species^4^. Critically, an active genetic element includes a guide RNA (gRNA) and is inserted into the genome at the precise location that is targeted for cleavage by the encoded gRNA. In a heterozygous animal that also expresses the Cas9 nuclease, the gRNA targets cleavage of the wild type homologous chromosome. Homologous sequences that flank the active genetic element then serve to resolve the double strand break by homology directed repair, which copies the active genetic element and converts the heterozygous genotype to homozygosity.

Despite the high efficiency observed in insects, the feasibility of CRISPR/Cas9-mediated active genetics has not yet been demonstrated in mammals. Divergence over 790 million years since the last common ancestor presents two potential obstacles to implementation in rodents: the efficiency of double strand break (DSB) formation using genetically encoded Cas9 and gRNA and/or the prevalence of homology directed repair may preclude efficient genotype conversion. Here, we report the feasibility of active genetics in laboratory mice.

Although homology directed repair (HDR) of CRISPR/Cas9 induced DSBs does occur *in vitro* and *in vivo* in mammalian cells and embryos, usually from a plasmid or single stranded DNA template, non-homologous end joining (NHEJ) is the predominant mechanism of DSB repair in somatic cells^5,6^. NHEJ frequently generates small insertions or deletions, known as “indels”, that make CRISPR/Cas9 an effective means of mutating specific sites in the genome. However, the competing pathway of homologous chromosome-templated HDR also has been shown to repair CRISPR/Cas9 induced DSBs in early mouse and human zygotes^7,8^. These observations raised the possibility that super-Mendelian inheritance of active genetic elements might also be achieved in mice.

To test this hypothesis, we designed a “CopyCat” element that differs from the self-propagating element used initially in insects in that it encodes a gRNA but does not encode the Cas9 protein, an arrangement that was also previously implemented in *Drosophila*^9^. We designed our strategy to disrupt the *Tyrosinase* gene (*Tyr*) because of the albino phenotype of homozygous loss-of-function animals^10^ and to make use of a previously characterized *Tyr* gRNA with high activity^11^. Figure 1a illustrates the insertion of this element precisely into the gRNA cut site in exon 4 of *Tyr* to obtain the *Tyr^CopyCat^* knock-in allele (Fig. 1a, Extended Data Fig. 1, 2, and Methods). Briefly, the *Tyr* gRNA is transcribed from a constitutive human RNA polymerase III U6 promoter^12^. On the reverse strand of DNA, to avoid possible transcriptional conflict, mCherry is ubiquitously expressed from the human cytomegalovirus early immediate promoter and enhancer (CMV)^13^. Since the 2.8 kb insert disrupts the *Tyr* open reading frame, *Tyr^CopyCat^* is a functionally null allele that is propagated by Mendelian inheritance in absence of Cas9. Crossing mice that carry the *Tyr^CopyCat^* element to transgenic mice that produce Cas9 allows us to test whether it is possible to observe super-Mendelian inheritance of the *Tyr^CopyCat^* allele (Fig. 1b). For this analysis, we took advantage of existing tools that provide spatial and temporal control of Cas9 expression and assessed eight different genetic strategies that vary CRISPR/Cas9 activity in the early embryo and in the male and female germlines.

**Figure 1.**
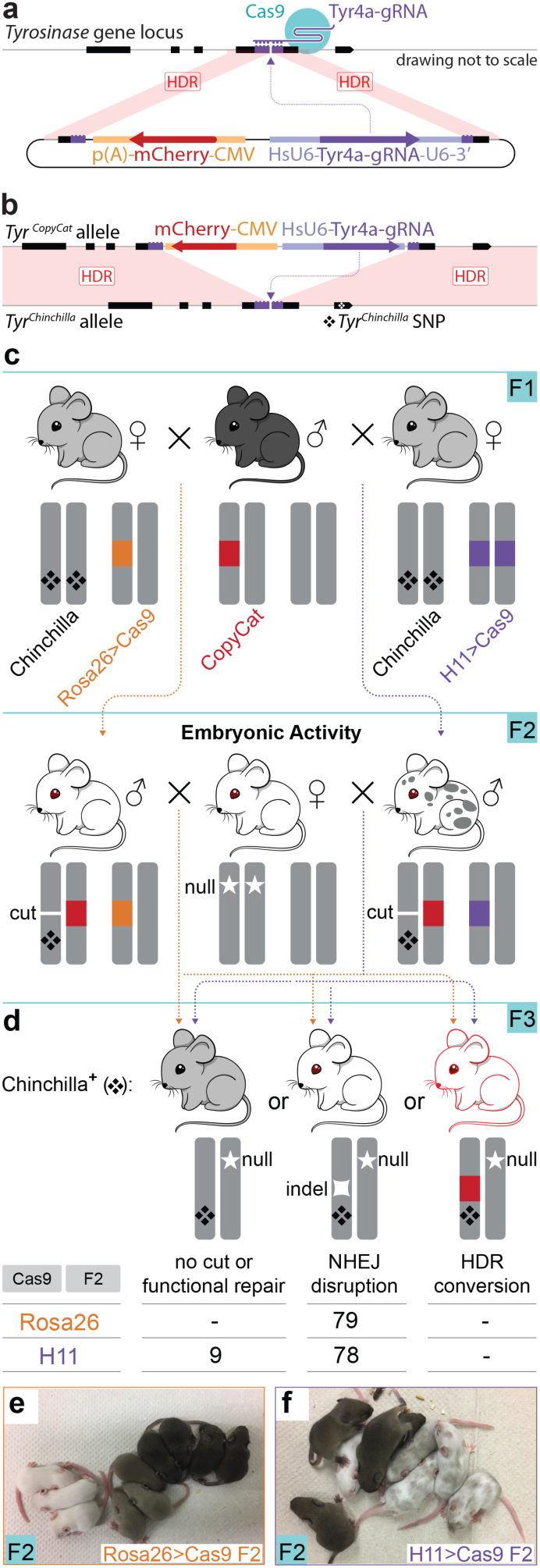
Embryonic Cas9 activity does not copy the *Tyrosinase^CopyCat^* allele from the donor to the receiver chromosome. (**a**) Knock-in strategy using the *Tyr^CopyCat^* targeting vector. The U6-Tyr4a gRNA and CMV-mCherry were inserted by HDR into the cut site of the Tyr4a gRNA. (**b**) The genetically encoded *Tyr^CopyCat^* element, when combined with a transgenic source of Cas9 is expected to induce a DSB in the *Tyr^Chinchilla^*-marked receiver chromosome, which could be repaired by inter-homologue HDR. (**c**) Breeding strategy to unite *Tyr^CopyCat^* with a constitutive *Cas9* transgene followed by test cross to *Tyr^Null^*. (**d**) Quantification of F3 test cross offspring. (**e**) A representative *Rosa26-Cas9* F2 litter. Black mice did not inherit *Tyr^CopyCat^*. Grey mice inherited *Tyr^CopyCat^* but not *Cas9*. White mice inherited both transgenes. (**f**) A representative litter in which all inherited *H11-Cas9*. The mosaic mice also inherited *Tyr^CopyCat^*.

We employed two “constitutive” *Cas9* transgenic lines, *Rosa26-Cas9*^14^ and *H11-Cas9*^15^, that reportedly express Cas9 in all tissues that have been assessed. Each knock-in allele is driven by a ubiquitous promoter and is placed in the respective *Rosa26*^16^ or *H11*^17^ “safe harbor” locus, which are thought to be accessible to the transcriptional machinery in all cells and thereby permit ubiquitous transgene expression. In order to track inheritance of the chromosome that is targeted for genotype conversion, we bred the *Chinchilla* allele of *Tyrosinase* (*Tyr^Ch^*)^18,19^ into each *Cas9* transgenic line to place a tightly linked genetic marker on the receiver chromosome. *Tyr^Ch^* encodes a hypomorphic point mutation in exon 5, and homozygotes or heterozygotes complemented with a null allele have a grey coat color^18,19^. The G to A single nucleotide polymorphism can also be scored with certainty by PCR and DNA sequencing (Extended Data Fig. 4).

Female *Rosa26-Cas9; Tyr^Ch/Ch^* and *H11-Cas9; Tyr^Ch/Ch^* mice were each crossed to *Tyr^CopyCat/+^* males with the goal of uniting the gRNA and Cas9 protein in the early embryo (Fig. 1c). In absence of a second loss-of-function mutation in exon 4 of the receiver chromosome, *Tyr^CopyCat/Ch^* mice should appear grey (*Tyr^CopyCat/Ch^*; *Cas9*-mice in Fig. 1e). However, we did not observe any grey *Tyr^CopyCat/Ch^; Cas9+* mice in the F2 offspring of either cross. Instead, 94% of *Rosa26-Cas9*; *Tyr^CopyCat/Ch^* mice were entirely white (17 white: 1 mosaic). In the case of *H11-Cas9; Tyr^CopyCat/Ch^* mice, 87.5% of the F2 progeny were a mosaic mixture of grey and white fur (21 mosaic: 3 white) (Fig. 1e, f and Supplementary Table 4). These results illustrate the highly efficient action of the *Tyr* gRNA, as well as a qualitative difference in the efficiency and/or timing of Cas9 activity driven by the *Rosa26-Cas9* versus the *H11-Cas9* transgenes.

Our next goal was to determine what type of repair events (NHEJ mutations or gene conversions by inter-homologue HDR) were transmitted to the next generation. In the absence of gene conversion, the expectation is that the *Tyr^CopyCat^* allele would be transmitted by Mendelian inheritance to 50% of the progeny of an outcross. In such cases, effectively none of the *Tyr^Ch-^*marked receiver chromosomes would be predicted to carry the *Tyr^CopyCat^* allele due to ultra-tight linkage of *Tyr* exons 4 and 5, which are separated by only ~9 kb. In order to assess inheritance in a large number of offspring, we crossed each F2 male *Rosa26-Cas9*; *Tyr^CopyCat/Ch^* and *H11-Cas9*; *Tyr^CopyCat/Ch^* mouse to multiple albino CD-1 females (*Tyr^Null^*), which carry a loss-of-function point mutation in *Tyr* exon 1^10,19^ (Fig. 1d). We then genotyped F3 offspring of this test cross by PCR and DNA sequencing to identify offspring that inherited the *Tyr^Ch^*-marked receiver chromosome.

In absence of a second null mutation induced in exon 4 of the *Tyr^Ch^*-marked receiver chromosome, *Tyr^Ch/Null^* animals should appear grey due to partial activity of the hypomorphic *Tyr^Ch^* allele. However, all F3 offspring of the *Rosa26-Cas9* lineage and 89.7% of F3 offspring of the *H11-Cas9* lineage were white, indicating the transmission of a CRISPR/Cas9 induced loss-of-function mutation on the receiver chromosome. The high frequency of mutation of the receiver chromosome in F3 offspring is consistent with the primarily albino coat color of the F2 parents (Fig. 1d, e, and f and Supplementary Table 4 and 5). If the induced null alleles resulted from inter-homologue HDR to copy the *Tyr^CopyCat^* allele from the donor to the *Tyr^Ch^*-marked receiver chromosome, these white animals should also express the fluorescent mCherry marker. In these experiments, however, none of the F3 offspring that inherited the receiver chromosome in either the *Rosa26*- or *H11-Cas9* lineages expressed mCherry. Confirming the lack of mCherry expression, PCR amplification of *Tyr* exon 4 revealed NHEJ-induced indels in white progeny (Fig. 1d and Extended Data Fig. 5 and 6).

The different propensities to yield full albino versus mosaic coat color patterns in the *Rosa26-Cas9* and *H11-Cas9* lineages were also paralleled by differences in the number of unique NHEJ mutations we identified on receiver chromosomes in individuals of each genotype. Sequenced PCR products from *Rosa26-Cas9; Tyr^CopyCat^* F2 tails (somatic tissues comprised of both ectodermal and mesodermal derivatives) and from individual F3 outcross offspring (representing the germline) routinely exhibited only two unique NHEJ mutations (Extended Data Fig. 6) suggesting that many of these Cas9 induced mutations were generated in 2-4 cell stage embryos. In contrast, *H11-Cas9; Tyr^CopyCat^* F2 tails and F3 offspring harbored several different NHEJ mutations, which suggests that Cas9 is active at later embryonic stages and/or at lower levels in this lineage (Extended Data Fig. 6).

The formation of indels in the early embryo provides an efficient method to generate mutations in a given gene with a low level of mosaicism to produce predictable whole organism phenotypes. Since such mutations are generated with high efficiency using *Rosa26-Cas9* transgenic mice, it should be possible to design an active genetic element encoding several gRNAs that target multiple genes simultaneously to evaluate the consequence of combinatorial gene knock-outs in a simple heritable system. These results are also relevant to recent reports showing that early zygotic CRISPR/Cas9 induced DSBs are repaired by inter-homologue HDR in mouse and human embryos^7,8^. We hypothesize that CRISPR/Cas9 activity and/or gRNA expression might be initiated too late in our genetic strategy to engage meiotic HDR mechanisms that were proposed to persist into the early zygote. The presence of so few unique NHEJ mutations in the *Rosa26-Cas9* lineage, however, suggests that zygotic inter-homologue HDR is transiently limited to a window of time very near fertilization.

We considered two reasons for the absence of *Tyr^CopyCat^* copying to the receiver chromosome in the early embryo. The first possibility is that homologous chromosomes are not aligned to allow for efficient strand invasion that is necessary for inter-homologue HDR to repair DSBs. Alternatively, the DNA repair machinery that is active in somatic cells typically favors NHEJ over HDR, which would generate indels that obliterate the gRNA cut site in the early embryo. A possible solution to overcome these two potential obstacles is to restrict CRISPR/Cas9 activity to meiosis during development of the germline. One purpose of sexual reproduction is to “shuffle the deck” by recombining the maternal and paternal genomes at each generation. Meiotic recombination is initiated by the intentional formation of DSBs that are repaired by exchange of DNA sequence information between homologous chromosomes that are physically paired during Meiosis I^20^. Indeed, the molecular mechanisms of NHEJ are repressed during meiosis in many species, including mice^21^, likely because activity of the NHEJ pathway in the germline would be highly mutagenic ^22^.

In order to test the hypothesis that Cas9 activity during meiotic recombination will convert a heterozygous active genetic element to homozygosity, we designed a crossing scheme to introduce the first expression of Cas9 in the presence of the *Tyr^CopyCat^* allele during germline development. Since there currently are no available transgenic mice that express Cas9 under direct control of a germline-specific promoter, we combined conditional *Rosa26*- or *H11-LSLCas9* transgenes, each with a Lox-Stop-Lox preceding the Cas9 translation start site^14,15^, with available *Vasa-Cre* or *Stra8-Cre* germline transgenic mice. *Vasa-Cre* is expressed later than the endogenous *Vasa* transcript in both male and female germ cells^23^ while *Stra8-Cre* expression is limited to the male germline and is initiated in early stage spermatogonia^24^. Although oogonia and spermatogonia are pre-meiotic, and spermatogonia are in fact mitotic, we reasoned that Cre protein must first accumulate to recombine the conditional *Cas9* allele for subsequent Cas9 protein expression and activity. The time delay may require initiation of *Cre* expression prior to the onset of meiosis so that Cas9-induced DSBs can be resolved by inter-homologue HDR prior to segregation of homologous chromosomes at the end of Meiosis I. We generated each combination of these *Cre* and conditional *Cas9* lines in case the timing or levels of Cas9 expression are critical variables in these crosses. We also assessed males and females of the *Vasa* strategies in case there are sex-dependent differences in animals that inherit the same genotype.

Males heterozygous for *Tyr^CopyCat^* and the *Vasa-Cre* transgene were crossed to females homozygous for both the *Tyr^Ch^* allele and one of the two conditional *Cas9* transgenes (Fig. 2a). We reasoned that in the reverse cross (i.e., using female *Vasa-Cre* mice), Cre protein that is maternally deposited in the egg might prematurely induce recombination of the conditional *Cas9* allele^23^. Early embryonic *Cas9* expression would have led to somatic mutagenesis similar to that observed in the experiments above using constitutive *Cas9* transgenes. Introducing the *Vasa-Cre* transgene by inheritance from the male resulted in most offspring that were entirely grey, due to the *Tyr^CopyCat/Ch^* genotype, and a few mosaic animals (Supplementary Table 6). The presence of any mosaicism suggests that even this approach for conditional germline restricted *Cas9* expression resulted in some degree of leakiness in somatic tissues.

**Figure 2.**
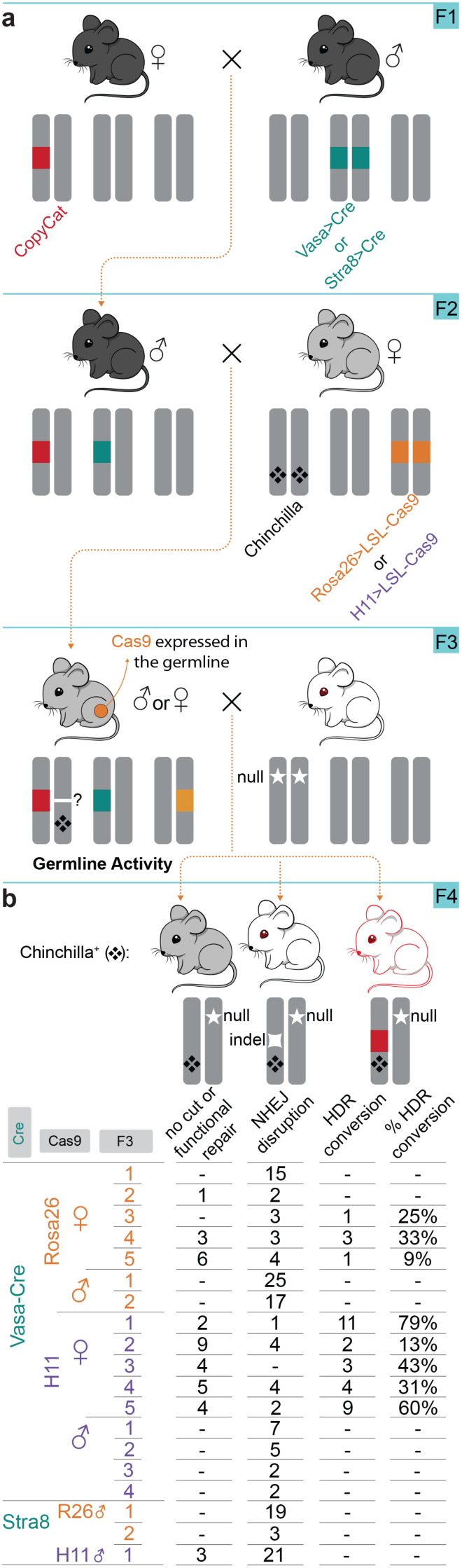
Cas9 activity in the female germline copies the *Tyrosinase^CopyCat^* allele from the donor to the receiver chromosome. (**a**) Breeding strategy to produce *Tyr^CopyCat/Chinchilla^* mice with a conditional *Cas9* transgene and a germline restricted *Cre* transgene. F3 offspring were test crossed to *Tyr^Null^* animals to assess F4 phenotypes and genotypes. (**b**) Quantification of the efficiency of HDR conversion in F4 test cross offspring.

We first tested whether expression of CRISPR/Cas9 in the female germline could promote copying of the *Tyr^CopyCat^* element onto the receiver chromosome by crossing female mice of each *Vasa-Cre* lineage to CD-1 (*Tyr^Null^*) males. In each cross, we identified offspring that inherited the *Tyr^Ch^*-marked chromosome. As in the cross to assess the effects of embryonic *Cas9* expression above, we expected *Tyr^Ch/Null^* mice without a second loss-of-function mutation in exon 4 of the receiver chromosome would be grey. Mice with a CRISPR/Cas9 induced NHEJ mutation in exon 4 would be expected to appear white. Mice with a CRISPR/Cas9 induced mutation that was repaired by inter-homologue HDR should also be white but additionally fluoresce red due to transmission of the *mCherry*-marked *Tyr^CopyCat^* active genetic element (Fig. 2b).

Figure 2b summarizes the results of these crosses that demonstrate genotype conversion upon Cas9 expression in the female germline. In contrast with constitutive embryonic expression of *Cas9*, we observed that the *Tyr^CopyCat^* transgene was copied to the *Tyr^Ch^*-marked receiver chromosome in both *Vasa-Cre; Rosa26-LSL-Cas9* and *Vasa-Cre;H11-LSL-Cas9* lineages. However, the observed efficiency differed between genotypes; three out of five females of the *Vasa-Cre;Rosa26-LSL-Cas9* lineage and all five of five females of the *Vasa-Cre;H11-LSL-Cas9* lineage transmitted a *Tyr^Ch^*-marked chromosome containing a *Tyr^CopyCat^* insertion to at least one offspring. Although there was considerable variation among females with the same genotype, the highest efficiency of genotype conversion within a germline produced 11 out of 14 *Tyr^Ch^* offspring (78.6%) with a *Tyr^CopyCat^* insertion in the *Vasa-Cre;H11-LSL-Cas9* lineage (Fig. 2b and Supplementary Table 7). These data demonstrate transmission of the *Tyr^CopyCat^* element that was copied onto the *Tyr^Ch^*-marked chromosome, an event with a very low probability (4.7×10^−5^) by natural recombination mechanisms due to tight linkage between the *Tyr^CopyCat^* and *Tyr^Ch^* alleles. Furthermore, although it seems likely that inter-homologue HDR of Cas9-induced DSBs utilizes the same DNA repair machinery that is active during meiotic recombination, these copying events cannot be explained by an increased incidence of chromosomal crossover, since all of the animals that inherited the donor chromosome that was not marked with *Tyr^Ch^* also expressed mCherry (Supplementary Table 7). In contrast, copying of the *Tyr^CopyCat^* element was not observed in any crosses where conditional *Cas9* expression was induced by *Vasa-Cre* in males. Consistent with this finding, *Tyr^CopyCat^* was also not copied to the *Tyr^Ch^*-marked receiver chromosome in the male germline of *Stra8-Cre* lineages (Fig. 2b and Supplementary Table 7).

These results reveal that active genetic elements can be copied to the homologous chromosome when Cas9 expression is restricted to the female germline, but these same strategies were unsuccessful in males. In mammals, spermatogonia continually undergo mitosis throughout the life of the male to produce new primary spermatocytes^25^. It is therefore possible that DSBs formed using the Cre-dependent strategies in males were repaired by NHEJ in mitotic spermatogonia, and the cut site was mutated prior to the onset of meiosis. In contrast, oogonia directly enlarge without further mitosis to form all of the primary oocytes^26^. These arrest during embryogenesis, prior to the first meiotic division, and oocyte maturation and meiosis continues after puberty. The higher efficiency of inter-homologue HDR in females of the *H11-LSL-Cas9* conditional strategy may reflect lower or delayed *Cas9* expression from the *H11* locus compared to *Rosa26*, also evident from a comparison of coat colors in the constitutive crosses. Thus, in the *Vasa-Cre;H11-LSL-Cas9* mice, Cas9 activity may have been fortuitously delayed even further to fall within an optimal window during female meiosis. The observed difference in the efficiency of inter-homologue HDR between females and males and even among females therefore likely indicates a requirement for the precise timing of CRISPR/Cas9 activity; NHEJ indels might reflect DSB repair that occurred prior to alignment of homologous chromosomes during Meiosis I or after their segregation (Fig. 3).

**Figure 3.**
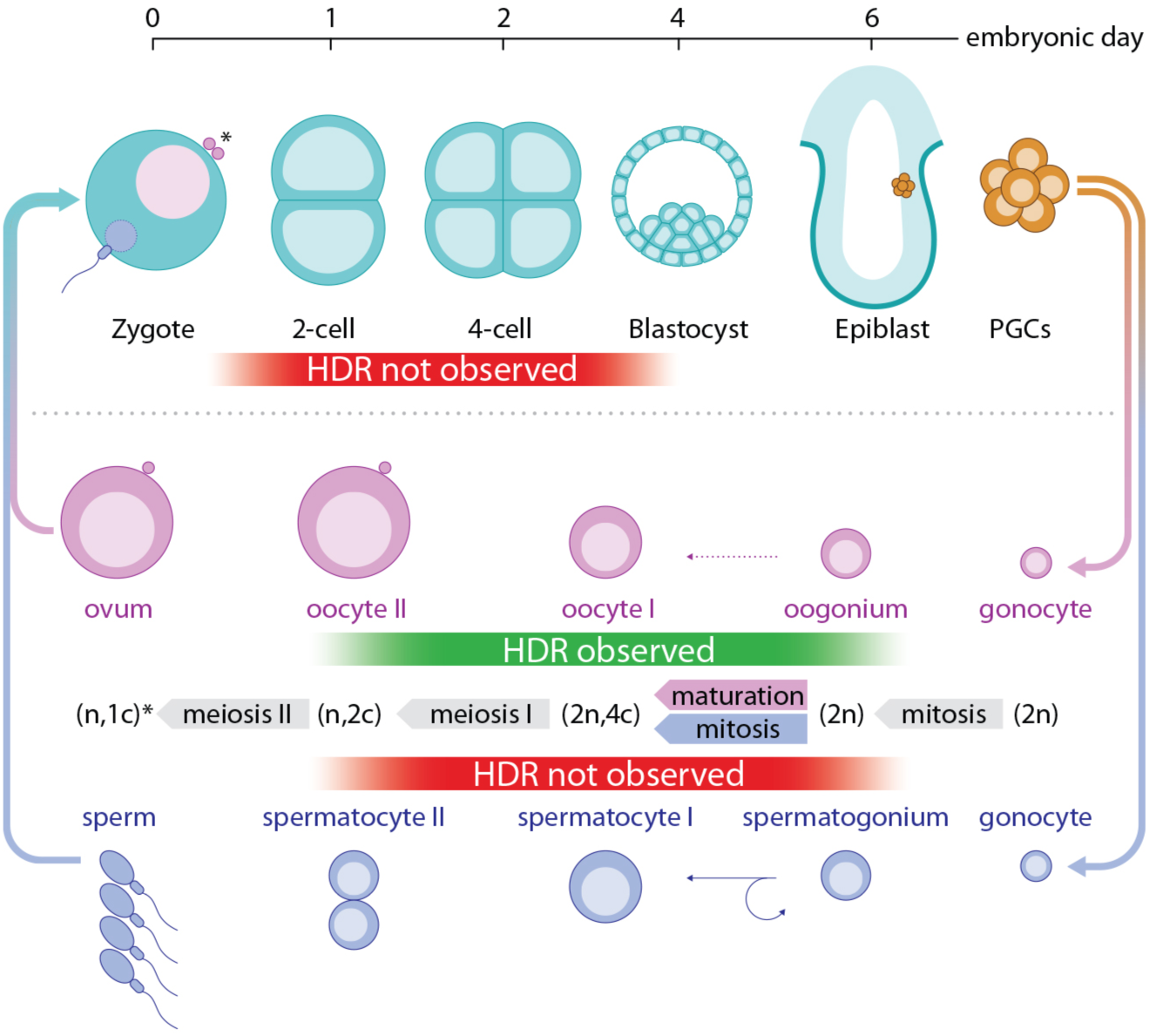
Genotype conversion by an active genetic element was observed in the female germline and not in the male germline or in the early embryo. Schematic representation of early embryonic and male and female germline development. Differences in germline specification coincide with presence or absence of observed HDR. [PGCs: primordial germ cells, n: number of homologous chromosomes, c: chromosome copy number. Asterisk indicates the difference between male sperm (n, 1c) and female ovum, which remains (n, 2c) until second polar body extrusion after fertilization.]

Our results comparing eight different genetic strategies indicate that precise timing of Cas9 activity may be more critical in rodents than in insects in order to leverage the endogenous meiotic recombination machinery for highly efficient gene drive. It therefore appears that both the optimism and concern that gene drives may soon be used to reduce invasive rodent populations in the wild is likely premature. These data are an important contribution to that ongoing discussion and to the prospects for further optimization. Nevertheless, while currently inadequate for gene drive implementation in the wild, the efficiency we demonstrate here is more than sufficient for laboratory applications and represents a significant advance in this arena. For example, the average copying rate of 45% using the most efficient genetic strategy (*Vasa-Cre;H11-LSL-Cas9*) could be used to combine even ultra-tightly linked loci such that approximately 23% of all offspring would inherit a chromosome with both alleles, an impossibility by Mendelian inheritance. This average copying rate also increases the probability of inheritance of a single desired allele from 50% to 73%. The highest rate of genotype conversion we observed (79%) would result in a 90% frequency of transmitting a desired allele. Such high transmission frequencies that both bypass the onerous constraint of genetic linkage and that increase the probability of obtaining animals homozygous for multiple genes should greatly accelerate the production of complex mammalian genotypes. Although there is opportunity to further improve the efficiency in the female and male germline, our results demonstrate a substantial advance that decreases the cost, time, and animal requirements for complex mammalian genetics.

## Acknowledgments

We thank Angela Green and An-Chih Chen for assistance with genotyping, Kaisa Hanley for providing the DNA extraction protocol, and Dr. Prashant Jain for assistance with fibroblast transfection to assess mCherry fluorescence from the *Tyr^CopyCat^* construct prior to transgenesis. We also thank Dr. Heidi Cook-Andersen and Dr. Miles Wilkinson for insightful conversations about mouse germline development and Dr. Luis Montoliu for discussion of the *Tyrosinase* locus and previously identified mutations. We thank Dr. Mark Tsuzynski for plasmid reagents and early support of the project. Expert animal care was provided by the UC San Diego Animal Care Program staff.

## Funding

This work was supported by a Searle Scholar Award from the Kinship Foundation, a Pew Biomedical Scholar Award from the Pew Charitable Trusts, and a Packard Fellowship in Science and Engineering from the David and Lucile Packard Foundation awarded to KLC. EB was supported by NIH grant R01 GM117321, a Paul G. Allen Frontiers Group Distinguished Investigators Award, and a generous gift from the Tata Trusts in India to TIGS-UCSD and TIGS-India. HAG was supported by a Ruth Stern Graduate Fellowship and by the NIH Cell and Molecular Genetics training grant T32 GM 7240-39A1; VMG was supported by the NIH DP5 OD023098.

## Author contributions

Conceptualization, HAG, VMG, GP, EB, and KLC; Resources, HAG, VMG, GP; Investigation, HAG, XSX; Data Curation, HAG; Formal Analysis, HAG; Supervision, KLC and EB; Writing-Original Draft, KLC; Writing – Review & Editing, HAG, VMG, EB, KLC.

## Competing interests

VMG, EB, and KLC hold advisory board positions with Synbal, Inc. All other authors declare that they have no competing interests.

## Data and materials availability

All data is available in the main text or the supplementary materials.

## Supplementary Materials

Methods

Extended Data Figures 1-6

Supplementary Tables 1-7

